# Daily membrane capacitance changes in mouse neurons

**DOI:** 10.1101/2022.12.09.519806

**Authors:** Daniel Severin, Cristián Moreno, Trinh Tran, Christian Wesselborg, Sofia Shirley, Alfredo Kirkwood, Jorge Golowasch

## Abstract

Capacitance of biological membranes is determined by the properties of the lipid portion of the membrane, as well as morphological features of a cell. In neurons, membrane capacitance is a determining factor of synaptic integration, action potential propagation speed and firing frequency due to its direct effect on the membrane time constant. Besides slow changes associated with increased morphological complexity during postnatal maturation, neuron membrane capacity is largely considered a stable, non-regulated constant magnitude. Here we report that in two excitatory neuronal cell types, pyramidal cells of mouse primary visual cortex and granule cells of the hippocampus, the membrane capacitance significantly changes between the start and the end of a daily light cycle. The changes are large, nearly two-fold in magnitude in pyramidal cells, but are not observed in cortical parvalbumin-expressing inhibitory interneurons. We discuss potential functional implications and plausible mechanisms.

## Introduction

The capacitance is a so-called passive property of the membrane and is a determinant factor of fundamental neuronal properties, such as action potential propagation speed (Hodgkin and Huxley 1952), synaptic integration (Baufreton et al. 2005; Martin 1976) and action potential firing frequency (Tewari et al. 2018). As summarized in equation [1],

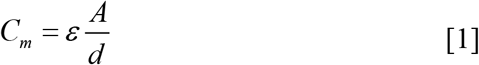

capacitance depends not only on the dielectric constant (*ε*) of the insulating material (phospholipid bilayer in biological membranes), but also on the distance between the intracellular and the extracellular conductive milieu (*d*), and the membrane area (*A*). Membrane capacitance, *C*_*m*,_ changes during early postnatal development. In some neuronal types the capacitance becomes smaller as *d* increases due to myelin wrapping (Castelfranco and Hartline 2015) or perineuronal net (PNN) encapsulation (Tewari et al. 2018). On the other hand, in most neurons total membrane capacitance increases as the membrane area increases during the maturation of structural complexity. It is common practice to report changes in capacitance as a proxy for changes in membrane surface (Haedo and Golowasch 2006; Perez-Garcia et al. 2021; Pineda et al. 2008; Royeck et al. 2008). In juveniles and adults, altered capacitance has only being reported for some mouse models of some neurological conditions or after traumatic insult (Akopian et al. 2016; Amzica and Neckelmann 1999; Perez et al. 2021; Rangel-Barajas et al. 2021; Tewari et al. 2018). Indeed, after maturation is completed, membrane capacitance is largely considered to be a constant biophysical property of the neurons and to be physiologically unregulated, since membrane thickness (*d*) and the properties of membrane lipids, captured by the dielectric constant (*ε*), are also thought to be stable (Gentet et al. 2000).

Here we report a 60-100% daily oscillation of membrane capacitance in excitatory pyramidal cells of mouse visual cortex from layers 2/3 and hippocampus granule cells. In contrast, parvalbumin-containing inhibitory interneurons (PV^+^) of the same visual cortex region show no such variations. We discuss the possible consequences for circuit stability, activity homeostasis and possible compensatory mechanisms that may operate simultaneously.

## Methods

### Animals

Both male and female animals were used. For visual cortex neuronal recordings we used 20-26 days old PV-cre;Ai14 mice, and for hippocampal neuronal recordings we used 114-128 days (∼4 months) old C57BL/6 mice (both from Jackson Laboratories, Bar Harbor Maine). All experiments were performed in accordance with the U.S. Public Health Service Policy on Humane Care and Use of Laboratory Animals, the National Institutes of Health Guidelines for the Care and Use of Animals in Research, and approved by the Institutional Animal Care and Use Committee at Johns Hopkins University, where the recordings were performed.

### Animal light entrainment and slice preparation

Mice were “entrained” for at least 2 weeks in light-controlled chambers with 12-hour light/dark cycles, scheduled to allow all experiments to be performed between 9 am and 8 pm. For slice preparation, the mice were removed from their cages ∼10 minutes before the chosen circadian time of study (*Zeitgeber*, ZT = 0/24, 6, 12 or 18 hours). The mice were first deeply anesthetized with isofluorane within 10 min after removal from their cages and then perfused transcardially with cold dissection buffer (5 ml at 10 ml/min) containing 212.7 mM sucrose, 5.0 mM KCl, 0.5 mM CaCl_2_,10 mM MgSO_4_ 1.25 mM NaH_2_PO_4_, 26 mM NaHCO_3_, and 10 mM glucose. After decapitation, brains were quickly removed, and acute sagittal brain ∼300 μm thick slices were made for visual cortex slices, and hippocampal slices (also ∼300 μm thick) were made as described in (Boric et al. 2008), both in ice-cold dissection buffer bubbled with a mixture of 5% CO_2_ and 95% O_2_. The slices were allowed to recover for 30 min at 30°C in dissection buffer and then for one hour at room temperature in artificial cerebrospinal fluid (ACSF): 124 mM NaCl, 5 mM KCl, 1.25 mM NaH_2_PO_4_, 26 mM NaHCO_3_, 1.5 mM MgCl_2_, 2.5 mM CaCl_2_, and 10 mM dextrose, and bubbled with a mixture of 5% CO_2_ and 95% O_2_.

All recordings were performed in a submerged recording chamber superfused with ACSF (30 ± 0.5°C, 2 ml/min). Synaptic blockers were included in the bath: 25 μM 6-cyano-7-nitroquinoxaline-2,3-dione (CNQX) to block AMPA/kainate receptors, 100 μM DL-2-amino-5 phosphonopentanoic acid (APV) to block NMDA receptors, and 10 μM gabazine to block GABA_A_ receptors. Whole-cell voltage-clamp recordings were obtained from pyramidal cells identified by their shape and firing response to depolarizing current pulses (Fig. 1) in cortical layers 2 and 3, and from identified parvalbumin (PV) cells (fluorescent in PV-cre;Ai14 mice). Hippocampal granule cells (GCs) were identified by their shape and position in the upper blade of the dentate gyrus (DG). We used borosilicate glass patch pipettes (3-6 MΩ) filled with intracellular solution containing the following: 130 mM K-gluconate, 10 mM KCl, 0.2 mM EGTA, 10 mM HEPES, 4 mM MgATP, 0.5 mM Na_3_GTP, 10 mM Na-phosphocreatine (pH 7.2–7.3, 280–290 mOsm). Average input resistance of pyramidal cells was 81.9 ± 21.7 MΩ (range: 64.4 to 115.0 MΩ), of PV cells was 148.7 ± 41.3 MΩ (range: 51.2 to 306.0 MΩ), and of granule cells was 249.2 ± 79.9 MΩ (range: 153.0 to 512.8.0 MΩ). Series resistance was < 20 MΩ (range 6-20MΩ), which was compensated at least 80% in every case. All drugs were purchased from either Sigma Aldridge (RRID:SCR_008988) or Tocris (RRID:SCR_003689).

**Figure 1.**
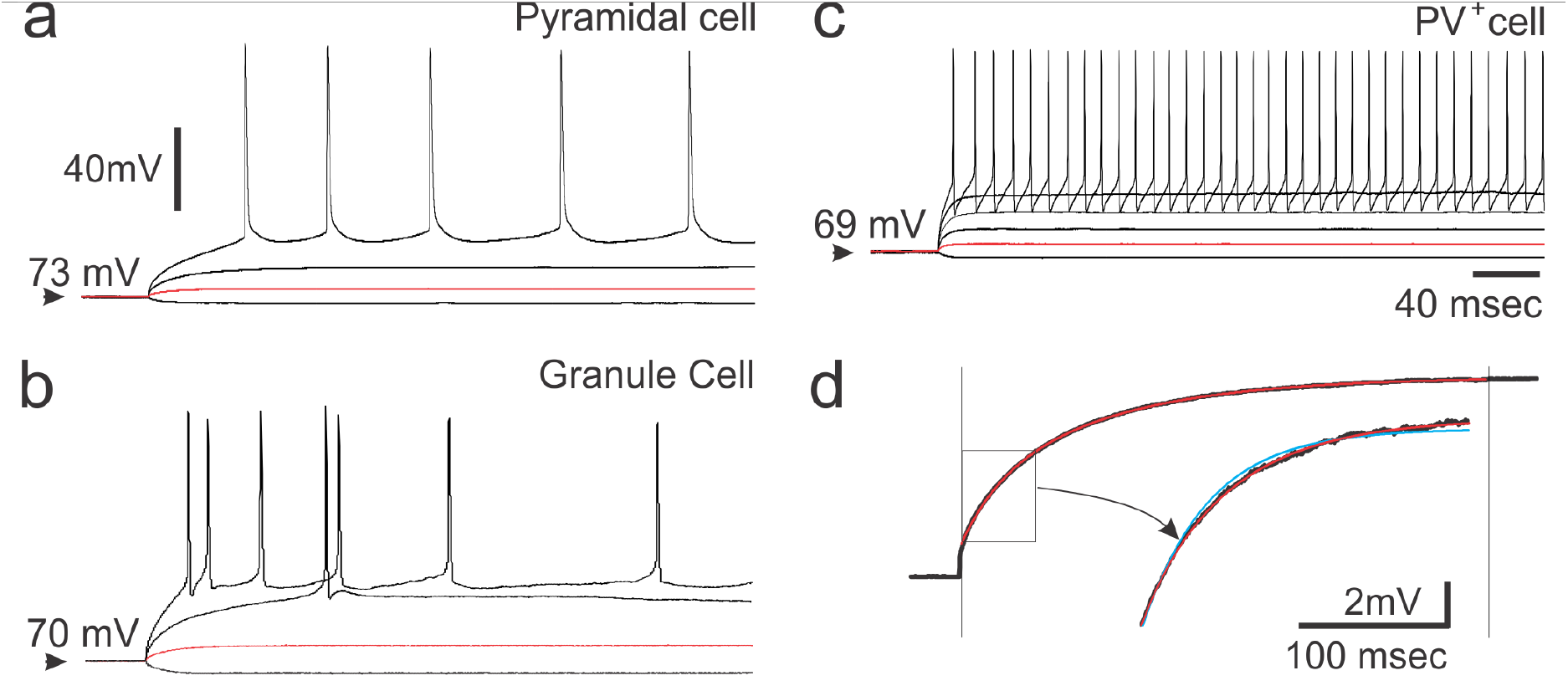
Typical neuronal activity of pyramidal cell, PV^+^cell and granule cell in response to current pulses. (**a**) Cortical pyramidal neurons and (**b**) hippocampal granule cells display a low frequency of action potential firing compared to (**c**) cortical PV^+^ cells. A range of current pulses was applied and subthreshold voltage response (red traces) in each case was fitted with a double exponential function from which membrane capacitance was derived as described in methods. (**d**) Example of a double exponential fit. The vertical lines indicate the limits for the 2-exponential fit (red trace) to a data trace (black). The boxed portion of the trace is shown amplified with a 2-exponential fit (red trace) and a single exponential fit (blue trace) superimposed to illustrate the improved goodness of the fit with two over one exponential.

### Capacitance measurements

Membrane capacitance was measured as described in Golowasch et al (2009). In that article it was demonstrated mathematically and computationally that in anisotropic cells, such as highly branched neurons, only membrane capacitance estimates from current clamp (and not from voltage clamp) experiments are a good approximation to their actual total membrane capacitance due to how the membrane potential of voltage clamped or current clamped cells behave in an electrotonically distributed cell. Using crab stomatogastric neurons, we also showed that in such cells voltage clamp estimates may underestimate membrane capacitance values by more than 10-fold in (Golowasch et al. 2009). Thus, here we used small depolarizing current pulses from a voltage around or slightly more depolarized than the resting potential of the cells (Resting potentials: Pyramidal cells: -70.2 ± 6.5 mV, N = 39; PV^+^ cells: -63.9 ± 5.2 mV, N = 45; GCs: - 64.0 ± 8.6 mV, N = 30). The rationale for the chosen baseline voltage and the pulse amplitude was to avoid or minimize entering a voltage range where voltage-dependent currents are activated during the pulse (White and Hooper 2013), thus remaining within levels where the cells respond passively. 1 second-long pulses were repeated at least 5 times, averaged to reduce noise and then baseline subtracted. It is important to avoid slow drifts or instabilities that may arise during very long pulses as they can strongly bias the passive membrane property measurements (White and Hooper 2013). We thus cut the length of the window to between the first 300-500 milliseconds to minimize slow drifts that may be observed in the trace and fitted the resulting trace with a double exponential function (Fig. 1d). The slowest exponential component corresponds to the cell’s membrane charging curve (Rall 1977), from which the time constant (*τ*_*m*_) can be obtained, and the total membrane capacitance (*C*_*m*_) and the total membrane resistance (*R*_*m*_) can be calculated (Golowasch et al. 2009; Rall 1977).

### Statistical Analysis

Figures show means and standard deviations, and tables show medians and quartiles. Data were statistically compared with one-way ANOVA tests for independent samples when normally distributed and non-normally distributed data were compared using Kruskal-Wallis ANOVA or Mann-Whitney Rank Sum *t*-test. These statistical analyses were performed using SigmaStat and graphs were made with SigmaPlot (Systat Software, Inc., San Jose, CA. USA; RRID:SCR_010285) and CorelDraw (Corel Inc., Austin, TX).

## Results

### Membrane capacitance varies as a function of time of day

We evaluated daily changes in the membrane capacitance, *C*_*m*_, in three types of identified neurons: pyramidal (Pyr) cells and parvalbumin positive inhibitory neurons (PV^+^) located in layers 2/3 of primary visual cortex, and granule cells (GC) from the hippocampus dentate gyrus. In each case, *C*_*m*_ was calculated from current clamp measurements of the membrane time constant, which was determined with multi-exponential fits to current-pulse induced voltage responses (see Methods, Fig. 1). We use this approach because it yields more accurate values of total membrane capacitance than estimations based on current responses to voltage clamp pulses. The commonly used voltage-clamp method significantly underestimates *C*_*m*_ due limitations imposed by the electrotonic structure of morphologically complex neurons (Golowasch et al. 2009). Surprisingly, we observed a nearly two-fold *C*_*m*_ decrease from the beginning of the light period (ZT0) to the beginning of the dark period (ZT12) in pyramidal cells (Fig. 2). In hippocampal granule cells we also we observed a large and significant *C*_*m*_ decrease (∼60%) between ZT0 and ZT12 (Fig. 2). These changes are highly statistically significant in both cell types (Fig. 2, Table 1).

**Table 1.**
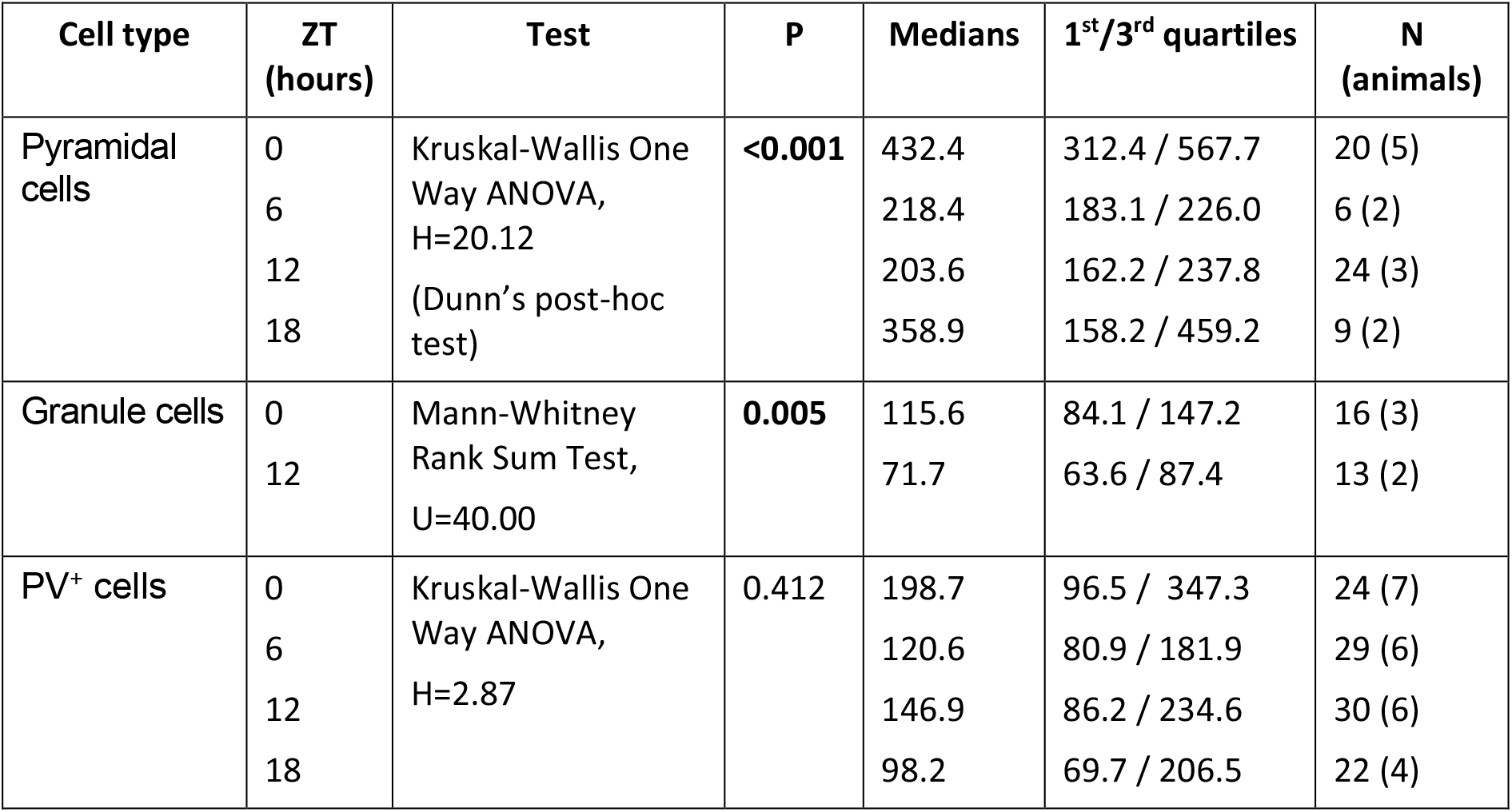
Statistics of comparisons of <u>membrane capacitance</u> at different times during one day (ZT) for cortical pyramidal and PV^+^ cell, and hippocampal granule cells. 1^st^ and 3^rd^ quartiles given. N = number of cells recorded (animals = number of animals from which these cells were taken).

| Cell type | ZT (hours) | Test | P | Medians | 1<sup>st</sup>/3<sup>rd</sup> quartiles | N (animals) |
| --- | --- | --- | --- | --- | --- | --- |
| Pyramidal cells | 0 | Kruskal-Wallis One Way ANOVA, H=20.12<br>(Dunn's post-hoc test) | <0.001 | 432.4 | 312.4 / 567.7 | 20 (5) |
|  | 6 |  |  | 218.4 | 183.1 / 226.0 | 6 (2) |
|  | 12 |  |  | 203.6 | 162.2 / 237.8 | 24 (3) |
|  | 18 |  |  | 358.9 | 158.2 / 459.2 | 9 (2) |
| Granule cells | 0 | Mann-Whitney Rank Sum Test, U=40.00 | 0.005 | 115.6 | 84.1 / 147.2 | 16 (3) |
|  | 12 |  |  | 71.7 | 63.6 / 87.4 | 13 (2) |
| PV<sup>+</sup> cells | 0 | Kruskal-Wallis One Way ANOVA, H=2.87 | 0.412 | 198.7 | 96.5 / 347.3 | 24 (7) |
|  | 6 |  |  | 120.6 | 80.9 / 181.9 | 29 (6) |
|  | 12 |  |  | 146.9 | 86.2 / 234.6 | 30 (6) |
|  | 18 |  |  | 98.2 | 69.7 / 206.5 | 22 (4) |

**Figure 2.**
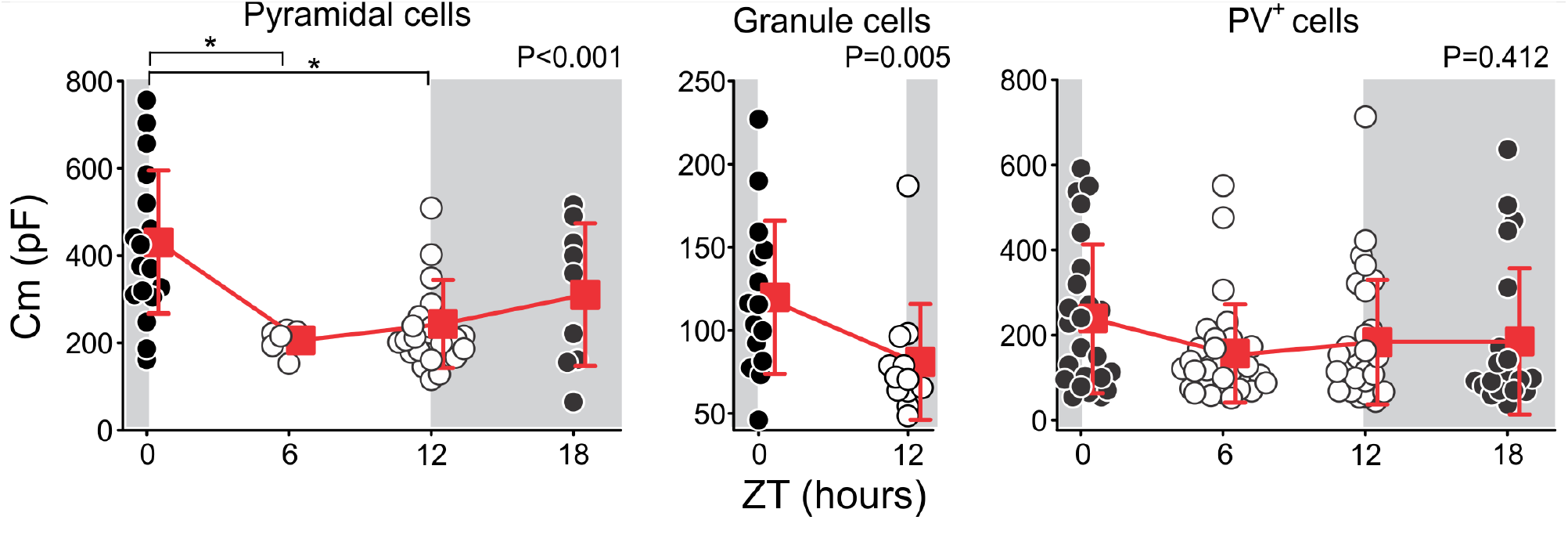
Daily membrane capacitance, C_m_, changes of cortical and hippocampal neurons. **Left.** *C*_*m*_ of visual cortex layer 2/3 pyramidal cells is highest at ZT0 (the end of the dark phase, gray area), and lowest at ZT6-ZT12 (the middle to end of the light phase, white area) (P < 0.001, Kruskal-Wallis One Way ANOVA). **Center**. Similar to pyramidal cells, *C*_*m*_ of hippocampus granule cells is highest at ZT0 and lowest at ZT12 (P = 0.005, Mann-Whitney Rank Sum Test). **Right.** *C*_*m*_ of cortical inhibitory PV^+^ neurons shows no variation over the course of a day (P = 0.412, Kruskal-Wallis One Way ANOVA). In these three panels individual cell *C*_*m*_ values are represented by hollow circles during the light phase, and dark symbols during the dark phase. Average ± standard deviations are shown as red squares and connecting lines, slightly displaced from the individual data for clarity. The horizontal brackets (top of pyramidal cells) indicate the times between which significant differences are observed in *post hoc* tests (P < 0.05, Dunn’s test). Statistical details are listed in Table 1.

Since pyramidal and granule cells are both excitatory, we asked if the total membrane capacitance of identified inhibitory cells might also vary as a function of the time-of-day. Previous work has shown small but significant daily changes in PNN wrapping around parvalbumin-positive (PV^+^) inhibitory cells in many brain areas, including cortex and hippocampus, that could result in membrane capacitance changes (Pantazopoulos et al. 2020). We recorded the time constant and calculated total *C*_*m*_, in identified inhibitory PV^+^ neurons from layer 2 and 3 of the visual cortex. By contrast to the excitatory neurons, cortical inhibitory PV^+^ neurons did not show any significant membrane capacitance change in the course of a 24-hour day/light cycle (Fig. 2, Table 1).

Interestingly, we observed a similar time-of-day dependence of the time constant, *τ*_*m*_, itself that closely tracks the membrane capacitance changes in both cortical pyramidal and hippocampal granule cells (Fig 3, left and middle). The changes of *τ*_*m*_ with the time-of-day were statistically significant (Table 2), with the main differences arising from differences between ZT0 – ZT12 and ZT6 - ZT18 (P < 0.05; Fig. 3, left) for pyramidal cells. However, as with *C*_*m*_, *τ*_*m*_ did not significantly vary in PV^+^ neurons (Fig. 3, right; Table 2).

**Table 2.** Statistics of comparisons of <u>membrane time constants</u> at different times during one day (ZT). Medians, 1^st^ and 3^rd^ quartiles given. N = number of cells recorded (animals = number of animals from which these cells were taken).

| Cell type | ZT (hours) | Test | P | Medians | 1<sup>st</sup>/3<sup>rd</sup> quartiles | N (animals) |
| --- | --- | --- | --- | --- | --- | --- |
| Pyramidal cells | 0 | One way ANOVA, F=5.80<br>(Holm-Sidak post hoc test) | 0.002 | 12.5 | 9.4 / 14.5 | 20 (5) |
|  | 6 |  |  | 9.5 | 6.9 / 16.3 | 6 (2) |
|  | 12 |  |  | 9.5 | 7.2 / 11.8 | 24 (3) |
|  | 18 |  |  | 9.9 | 8.0 / 13.9 | 9 (2) |
| Granule cells | 0 | Mann-Whitney Rank Sum Test, U=32.50 | 0.002 | 11.6 | 7.3 / 17.1 | 16 (3) |
|  | 12 |  |  | 5.9 | 5.4 / 7.1 | 13 (2) |
| PV<sup>+</sup> cells | 0 | Kruskal-Wallis One Way ANOVA, H=5.04<br>(Dunn's post hoc test) | 0.169 | 12.5 | 9.4 / 14.7 | 24 (7) |
|  | 6 |  |  | 9.5 | 6.9 / 16.3 | 29 (6) |
|  | 12 |  |  | 9.5 | 7.2 / 11.8 | 30 (6) |
|  | 18 |  |  | 9.9 | 8.0 / 13.9 | 22 (4) |

**Figure 3.**
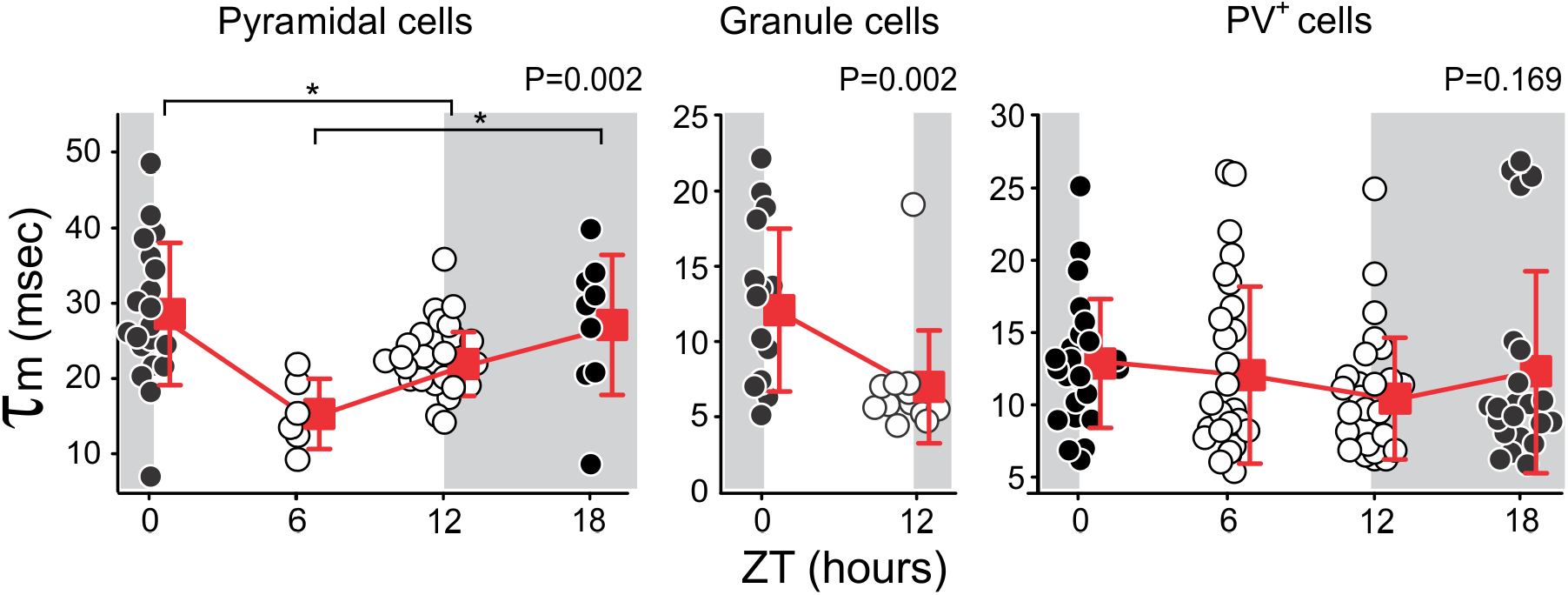
Daily membrane time constant, τ_m_, changes of cortical and hippocampal neurons. **Left.** *τ*_*m*_ of visual cortex layer 2/3 pyramidal cells is highest at ZT0 (end of the dark phase, gray area) and lowest at ZT6 (middle to end of the light phase, white area) (P < 0.002, One Way ANOVA). **Center.** *τ*_*m*_ of hippocampus granule cells is significantly higher at ZT0 than at ZT12 (P = 0.002, Mann-Whitney Rank Sum Test). **Right.** *τ*_*m*_ of cortical inhibitory PV^+^ neurons does not vary over the course of a day (P = 0.169, Kruskal-Wallis One Way ANOVA). Color of symbols and background have the same meaning as in Figure 1. The horizontal brackets (top of pyramidal cell data) indicate the times between which significant differences are observed in a *post hoc* test (P < 0.05, Dunn’s test). Statistical details are listed in Table 2.

Finally, *τ*_*m*_ is the product of the membrane capacitance and the membrane resistance, *R*_*m*_ (Eq. 1). Therefore, we calculated *R*_*m*_ in all three cell types to examine to what extent it may affect the temporal properties of the membrane in these cells. Figure 4 shows these data. We observed that only in pyramidal cells is there a significant effect of time-of-day on *R*_*m*_ (Fig. 4, left, Table 3). Both granule cells and PV^+^ cells show no significant *R*_*m*_ changes (Fig. 4, middle and right, Table 3). Importantly, the *R*_*m*_ changes observed in pyramidal cells run opposite to the changes in membrane time constant (Fig. 3, left), suggesting that *τ*_m_ changes are primarily due to membrane capacitance, and not membrane resistance, changes.

**Table 3.** Statistics of comparisons of <u>membrane resistances</u> at different times during one day (ZT). Medians, 1^st^ and 3^rd^ quartiles given. N = number of cells recorded (animals = number of animals from which these cells were taken).

| Cell type | ZT | Test | P | Medians | 1<sup>st</sup>/3<sup>rd</sup> quartiles | N (animals) |
| --- | --- | --- | --- | --- | --- | --- |
| Pyramidal cells | 0 | Kruskal-Wallis One Way ANOVA,<br>H=13.94<br>(Dunn's post-hoc test) | **0.003** | 68.4 | 58.2 / 87.5 | 20 (5) |
|  | 6 |  |  | 69.0 | 59.4 / 91.6 | 6 (2) |
|  | 12 |  |  | 112.0 | 99.9 / 145.8 | 24 (3) |
|  | 18 |  |  | 76.7 | 60.4 / 192.0 | 9 (2) |
| Granule cells | 0 | Mann-Whitney Rank Sum Test,<br>U=89.00 | 0.525 | 89.1 | 70.6 / 134.7 | 16 (3) |
|  | 12 |  |  | 84.0 | 77.0 / 96.5 | 13 (2) |
| PV<sup>+</sup> cells | 0 | Kruskal-Wallis One Way ANOVA,<br>H=4.86 | 0.182 | 59.9 | 41.3 / 109.2 | 24 (7) |
|  | 6 |  |  | 88.4 | 60.0 / 118.6 | 29 (6) |
|  | 12 |  |  | 69.2 | 48.6 / 91.9 | 30 (6) |
|  | 18 |  |  | 88.1 | 58.4 / 106.4 | 22 (4) |

**Figure 4.**
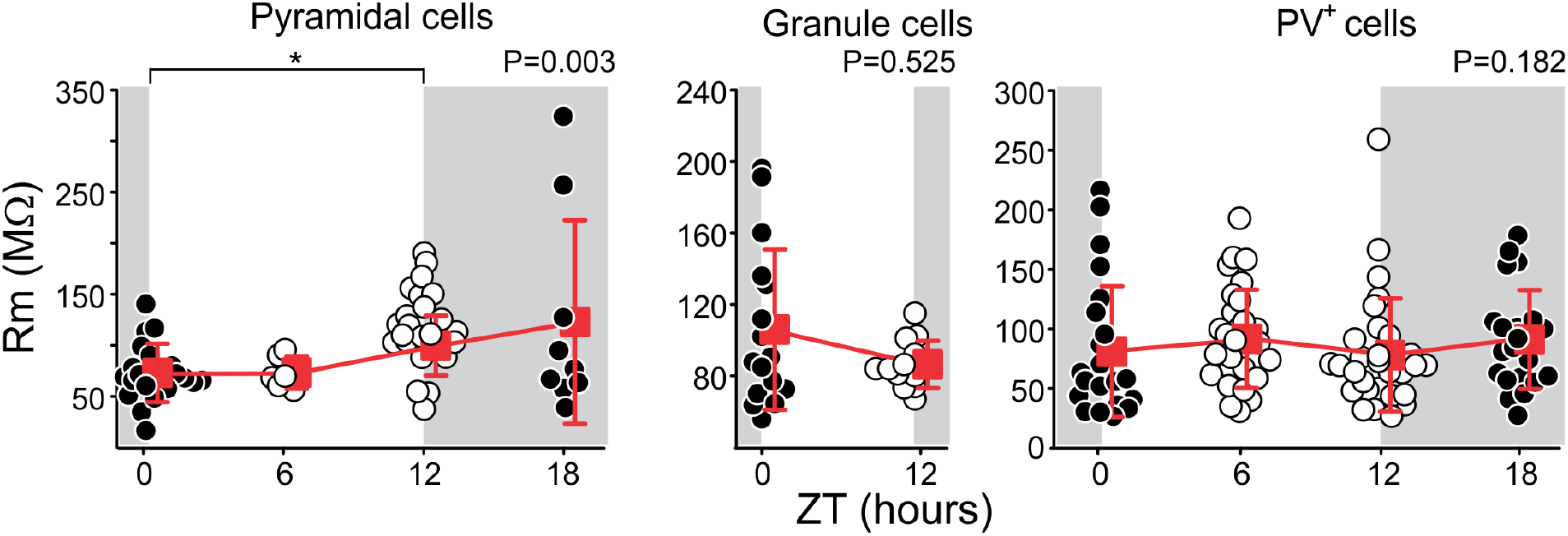
Daily membrane resistance, R_m_, changes of cortical and hippocampal neurons. **Left**. The membrane resistance of visual cortex layer 2/3 pyramidal cells is lowest at ZT0 (end of the dark phase, gray area) and ZT6 (middle of the light phase, white area) and highest at ZT12 (end of light phase) to ZT18 (middle of the dark phase) (P < 0.003, Kruskal-Wallis One Way ANOVA). **Center**. Membrane resistance of hippocampus granule cells does not change with time of day (P = 0.525, Mann-Whitney Rank Sum Test). **Right**. The membrane time constant of cortical inhibitory PV^+^ neurons also does not vary over the course of a day (P = 0.182, Kruskal-Wallis One Way ANOVA). Color of symbols and background have the same meaning as in Figure 1 and 2. The horizontal bracket (top of pyramidal cell data) indicates the times between which significant differences are observed in a *post hoc* test (P < 0.05, Dunn’s test). Statistical details are listed in Table 3.

## Discussion

Here we report that membrane capacitance exhibits a remarkably large daily oscillation (exceeding 60% change between peak and trough) in two types of excitatory neurons, visual cortex pyramidal cells and hippocampal granule cells. These changes occur with a maximum around ZT0 (end of the dark / beginning of the light period) and a minimum around ZT12 (end of the light / beginning of the dark period). In contrast, membrane capacitance did not change in visual cortical PV^+^ inhibitory neurons. Thus, in a manner highly reminiscent of our recent report of large daily changes in excitation/inhibition balance (Bridi et al. 2020), the capacitance changes are not restricted to a single brain region, yet within a region they do not occur in all neuron types. Together, both observations reveal an unexpected degree of complex neural remodeling in fundamental membrane properties previously considered stable. The amplitude of the membrane capacitance changes are surprisingly, exceeding by almost an order of magnitude membrane capacitance changes observed in previous studies (Akopian et al. 2016; Amzica and Neckelmann 1999; Perez et al. 2021; Rangel-Barajas et al. 2021; Tewari et al. 2018).

Multiple independent cellular mechanisms could potentially contribute to the daily modulation of capacitance. One candidate could be daily changes in membrane area resulting from the turnover of cortical synaptic spines, which are known to exhibit a circadian-like regulation in number and shape (de Vivo et al. 2017; Diering et al. 2017). These changes, however, affect a relatively small fraction of spines, and are thus highly unlikely to account for the 60-100% daily change in total membrane capacitance reported here. Similarly, the dendritic arborization of pyramidal neurons in the mouse visual cortex can increase substantially in length and complexity, but only for a short postnatal period in a process that is nearly completed by p21 in V1, which is the age of the youngest of our subjects (Richards et al. 2020). Furthermore, dendritic length and number changes as a function of visual experience are much smaller than the observed *C*_*m*_ changes (Richards et al. 2020).

Another possibility to consider is the recent report that conditioning artificial phospholipid membranes with voltage stimulation induces persistent, yet reversible, changes in the area and thickness of artificial bilayers, therefore affecting their capacitance (Scott et al. 2022). If this mechanism also operates in the brain, the changes in membrane capacitance would be expected to reflect daily oscillations in total neuronal activity.

A more plausible mechanism relates to changes in the extracellular space (ECS) volume. Consider a simple scenario in which current injected into a neuron sequentially crosses its own membrane, then an ECS of variable thickness and a layer of glial membrane to finally reach the reference electrode. In that case, the estimated *C*_*m*_ strongly depends on the ECS width as it would impact the effect membrane thickness (see Eq. 1). This scenario would predict local but extensive changes in ECS around cortical pyramidal and hippocampal granule cells, with ECS expanding during the light phase (when *C*_*m*_ drops) and shrinking during the dark phase (when *C*_*m*_ increases). The changes would need to be local since otherwise they would not explain the lack of *C*_*m*_ change in PV^+^ interneurons. Notably, another factor that affects membrane capacitance is the perineural network (PNN) that prominently encapsulates PV^+^ interneurons (Briones et al. 2021; Tewari et al. 2018; Wang and Fawcett 2012). One intriguing possibility is that the unique *C*_*m*_ stability in PV^+^ interneurons relate to their tight encapsulation by perineural networks (PNNs), which is not observed in either cortical pyramidal or hippocampal granule cells (Briones et al. 2021; Carceller et al. 2022; Tewari et al. 2018). This conjecture is built on two recent observations. On one hand, manipulations to reduce PNN density increase *C*_*m*_ in PV^+^ interneurons as expected from a thinning of the effective neuronal membrane thickness (Eq. 1) (Tewari 2018). On the other hand, PNN density on PV^+^ interneurons oscillates daily with a trough at ZT4-8 and a peak at ZT18-24 (Pantazopoulos et al. 2020). Thus, the same or a similar mechanism as the one acting on cortical pyramidal and granule cells to change *C*_*m*_ may also be acting on cortical PV^+^ interneurons, but with the PNN thickness around PV^+^ interneurons oscillating in phase, thus working in opposite directions on their *C*_*m*_. In such case, *C*_*m*_ oscillations would be noticeable only in cortical pyramidal and hippocampal granule cells but be dampened in PV^+^ interneurons.

Membrane capacitance changes can profoundly impact the physiology of the affected neurons and the networks they are part of through their observed effects on the membrane time constant. Three principal effects that changes in the time constant of a neuron can have are: changes in synaptic integration, action potential propagation speed and action potential spiking frequency. Synaptic integration is likely the most consequential of these effects because it plays a crucial role in the computation of inputs coming from different regions of the nervous system onto a target cell (Spruston 2008). Pyramidal cells integrate information from both top-down and bottom-up sources, with apical and basal dendrites handling these incoming signals, respectively. Similarly, granule cells receive inputs from the lateral and medial entorhinal cortices in the upper and lower portions of the perforant path, respectively. In addition, *C*_*m*_ changes will likely alter the velocity and reach of action potentials back-propagating into apical dendrites. In turn, this can affect the modification of distal synapses induced via spike-timing dependent plasticity (STDP), because STDP depends crucially on the coincidence of synaptic activation with the arrival of the action potential (see (Debanne and Inglebert 2023)).

The exact consequences of daily *C*_*m*_ changes in the function of actual neural circuits are not simple to predict because they likely occur in the context of multiple other daily neuronal changes such as those affecting ion conductance regulation (Adamsky et al. 2018), co-regulation of active conductance (Tran et al. 2019), and regulation of the excitation/inhibition balance (Bridi et al. 2020). Whether these various processes compensate or amplify the consequences of membrane capacitance changes remains to be determined. In either case, we propose that the observation of large daily membrane capacitance changes introduces a fundamental new variable to consider for understanding the dynamics of neuronal activity.

